# Unveiling the tempo of molecular and morphological evolution across the Tree of Life

**DOI:** 10.1101/2025.07.20.665814

**Authors:** E. Karen López-Estrada, Yasmin Asar, Hervé Sauquet, Simon Y. W. Ho

**Affiliations:** School of Life and Environmental Sciences, University of Sydney, NSW 2006, Australia; Unidad de Síntesis en Sistemática y Evolución, Instituto de Biología, Universidad Nacional Autónoma de México, 3er Circuito de Ciudad Universitaria, Coyoacán, Ciudad de México, 04510, Mexico; National Herbarium of New South Wales, Royal Botanic Gardens and Domain Trust, Sydney, NSW, Australia; Evolution and Ecology Research Centre, School of Biological, Earth and Environmental Sciences, University of New South Wales, Sydney, NSW 2052, Australia

**Keywords:** punctuated equilibria, phyletic gradualism, evolutionary rate, heterotachy, diversification

## Abstract

The evolution of Earth’s vast genetic and morphological diversity has been explained by an array of macroevolutionary models. At opposite ends of the spectrum lie two contrasting evolutionary models: phyletic gradualism and punctuated equilibria. Under a phyletic gradualism framework, evolutionary change accumulates along lineages and species are steadily transformed into new forms over time. In contrast, under punctuated equilibria, evolutionary change tends to occur in bursts at speciation events. Previous studies of molecular and morphological data have found varying levels of support for the two evolutionary models. We examined these models using comprehensive molecular and morphological data sets from 40 clades across the Tree of Life. Testing for associations between species richness and the amount of evolutionary change in sister clades, we find little evidence to support the punctuated equilibria model. However, we found high levels of among-lineage rate variation in molecular evolution and particularly morphological evolution. Our comparison of coding and non-coding genomic regions revealed contrasting patterns of among-lineage rate variation, without clear trends across taxa. Our study confirms that heterotachy is a dominant feature in macroevolution and that molecular and morphological evolution cannot simply be described by either a gradual or punctuated model.

## Introduction

The vast diversity across present-day organisms is the outcome of various evolutionary processes acting at different intensities and frequencies through time and among lineages. These heterogeneous patterns of macroevolutionary change reflect variation in both the *tempo* and *mode* of evolution across the Tree of Life (Harmon et al., 2003; Alfaro et al., 2009; Venditti et al., 2011). There have been many attempts to develop mechanistic and phenomenological models that can capture the key features of the macroevolutionary process, particularly with respect to the rate and timing of phenotypic and genomic change (e.g., Eldredge & Gould, 1972; Malmgren et al., 1983; Pagel et al., 2006; Douglas et al., 2025; Usai et al., 2024; Budd and Mann, 2025).

In the traditional Darwinian model of phyletic gradualism, evolutionary change occurs at a steady pace via the accumulation of incremental modifications (Darwin, 1859; Simpson, 1944). Macroevolutionary change is predominantly governed by natural selection and is interpreted to be an extension of microevolutionary processes across large spatial and temporal scales. Nevertheless, extensive analyses of molecular and morphological data have convincingly demonstrated that rate variation is a ubiquitous feature of evolution (e.g., May & Moore, 2019; Ho, 2020; Asar et al., in prep), prompting a need for macroevolutionary models that can account for shifts in rates across phylogenetic trees.

One prominent macroevolutionary model, punctuated equilibria (Eldredge & Gould, 1972), proposes that long periods of stasis are interrupted by bursts of evolutionary change concentrated at speciation events. Small population sizes, the release of evolutionary constraints, and “genetic revolutions ” are some of the mechanisms that have been put forward to account for these episodes of rapid evolutionary change (Mayr, 1954; 1963; Alberch, 1980; Venditti & Pagel, 2009). In response to evidence that such spikes of change did not always seem to occur at speciation events, subsequent models of punctuated gradualism or punctuated anagenesis proposed that pulses of change could occur at any time during the course of evolution, without necessarily being coincident with speciation events (Malmgren et al., 1983). Several methods have been developed to identify such shifts or pulses in morphological evolutionary rates, employing Lévy or compound Poisson processes to treat these pulses as rare events that can happen at any point along branches of the phylogeny (e.g., Landis et al., 2013; Rabosky et al., 2013; Pagel et al., 2022).

Analyses of morphological and molecular data have provided varying support for each of these macroevolutionary theoretical frameworks (Duchêne & Bromham, 2013; Rabosky et al., 2013; Kaji et al., 2018; López-Estrada et al., 2019; Heasly et al., 2021; Pagel et al., 2022; Brownstein et al., 2024; López-Martínez et al., 2024). In most cases, the findings were limited to specific groups of organisms or data types (e.g., only fossil taxa or only molecular data), but several broad-scale analyses concluded that punctuated change is an important feature of molecular evolution (Webster et al., 2003; Pagel et al., 2006; Douglas et al., 2025). Nevertheless, there has been persistent debate about the relative importance of gradual and punctuated modes of change (Gingerich, 1984; Von Vaupel Klein, 1995; Pennell et al., 2014), and it remains unclear whether such macroevolutionary patterns occur more broadly across phenotypic characters and genomes and throughout the Tree of Life.

The growing wealth of genomic and phenomic data presents valuable opportunities to test classic macroevolutionary models for a broad sample of organisms. Although new and more complex phenomenological models have been proposed to describe the timing and mode of evolutionary change (Usai et al., 2024; Budd and Mann, 2025; Douglas et al., 2025), the debate between gradual and punctuated evolution has been an enduring one (Heasley et al., 2021; Pagel et al., 2022). Here we test the model of punctuated equilibria by analysing a collection of 40 data sets, sampled from groups of organisms ranging from plants and insects to polychaetes and vertebrates. Using a sister-pairs approach to account for phylogenetic covariance and to mitigate the problems associated with node-density effects, we are able to compare and contrast the evolutionary patterns for morphological and molecular data across a large and diverse range of eukaryotes. By analysing large molecular data sets, we are able to distinguish between patterns in genomic regions under strong selection (ribosomal RNA genes, ultraconserved elements, and first and second codon sites of protein-coding genes) and weak selection (intergenic regions, introns, and third codon sites of protein-coding genes). Although we identify pronounced heterogeneity in evolutionary rates across lineages, our results collectively reveal a lack of compelling evidence to support the model of punctuated equilibria.

## Results and Discussion

### Evolutionary change linked to diversification is the exception not the rule

Our analyses of 40 data sets, representing a broad cross-section of the Tree of Life (Figure 1), reveal a lack of evidence for punctuated equilibria in morphological and molecular evolution. These results were obtained using phylogenetically independent contrasts or sister-pairs to test whether episodes of rapid evolutionary change are associated with cladogenetic events. For all 40 data sets that we analysed, we did not find a significant positive correlation between net diversification (measured by the number of extant species in a clade) and the amount of molecular or morphological change (measured by the phylogenetic branch length), regardless of taxonomic level, number of sister-pairs, or data set size. We did not detect any positive association even in the largest data sets, containing more than 40 independent contrasts and where a high level of statistical power would be expected. Our findings are broadly consistent with those of previous studies that found that punctuated equilibria explained only a small proportion of molecular evolutionary change (Webster et al., 2003; Pagel et al., 2006).

**Figure 1.**
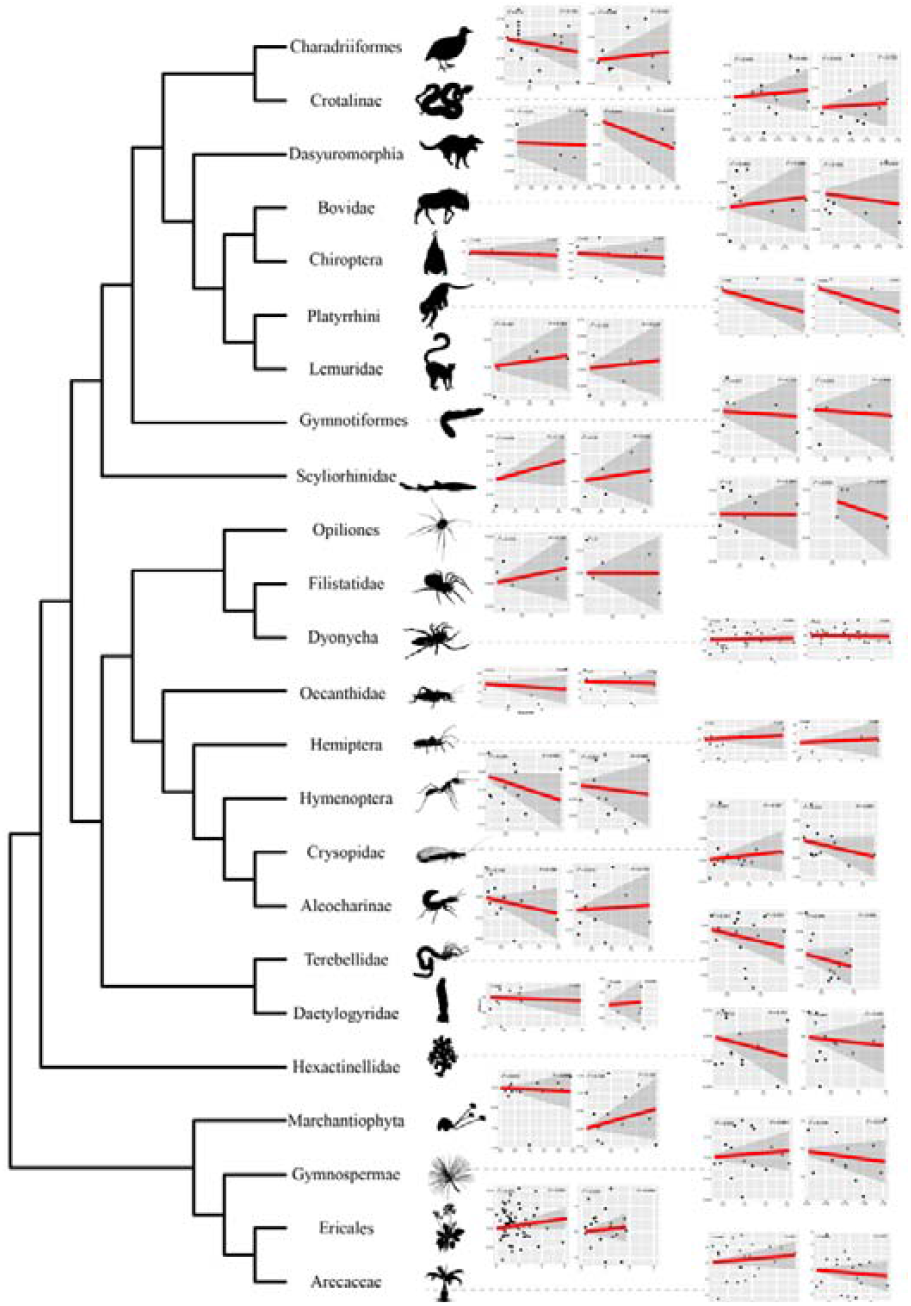
Tests of correlations between evolutionary rates and net diversification rates for 24 taxa across the eukaryotic Tree of Life, selected from 40 groups of taxa analysed in this study. Plots are shown for sister-pair contrasts in molecular rates (left panels) and contrasts in morphological rates (right panels). Lines of best fit are in red, with 95% confidence intervals in grey shading.

The lack of strong evidence of a link between the diversification process and the degree of morphological or molecular evolution is not entirely unexpected. Recent studies have shown that molecular and morphological evolutionary rates are largely uncoupled across the Tree of Life (Bromham et al. 2002; Asar et al. in prep). Furthermore, there is a growing catalogue of examples of lineages having diversified without any apparent morphological divergence (i.e., cryptic speciation; Wisecaver et al., 2023; Williamson et al., 2024; Grupstra et al., 2024). Although there are various reasons to believe that speciation can drive molecular evolution (Barraclough & Savolainen, 2001; Cooper & Purvis, 2009; Venditti & Pagel, 2009; Ritchie et al., 2022), our findings suggest that this association is either rare or difficult to detect.

In contrast with the predictions of punctuated equilibria, our analysis of data from shorebirds (order Charadriiformes) reveals a negative relationship between evolutionary rate in molecular data under strong selection and net diversification. This relationship is difficult to explain, given that greater genomic divergence would be expected to lead to reproductive incompatibilities that promote speciation (Dobzhanksy, 1937; Orr & Turelli 2001). However, such negative links between molecular evolutionary change and diversification have been reported in ferns and some groups of vertebrates (Testo & Sundue, 2018; Cooney & Thomas, 2021). Furthermore, previous studies have revealed cases of ‘static evolution’ (Janecka et al., 2012), where pronounced morphological or other forms of differentiation occur without corresponding molecular divergence; examples have been documented from primates and cichlids (King & Wilson, 1975; Rüber & Adams, 2001; Nakamura et al., 2021). In the case of Charadriiformes, we suggest that ecological or geographical differentiation, rather than morphological divergence, might have played a primary role in the diversification process, even though it is not accompanied by substantial molecular evolution. Nonetheless, our study reveals that a negative association between diversification and molecular change is an unusual case.

The lack of correlation between molecular evolutionary change in genomic regions under weak selection and net diversification might provide some insight into the dominant modes of genomic evolution. We have found that the amount of change in noncoding DNA is not tied to net diversification, in contrast with previous findings from birds (Lanfear et al. 2010b) and flowering plants (Duchêne & Bromham 2013). More broadly, our results are inconsistent with the prediction that a higher genome-wide mutation rate should lead to a greater rate of reproductive isolation (Orr & Turelli 2001), which should then increase the net diversification rate. Instead, our analyses of genome-wide data do not yield any evidence of a punctuational contribution at speciation events.

Finally, the absence of an association between evolutionary changes in coding DNA and net diversification may highlight the challenges of detecting punctuated equilibria as a mode of evolution. The punctuated equilibria model, grounded in the theory of allopatric speciation, posits that selection typically maintains equilibrium between populations and their environments (Eldredge & Gould, 1972). Consequently, adaptive changes or sites under strong selection are expected to exhibit clearer signatures of punctuated evolution. However, such adaptive signatures are usually limited to specific regions of the genome, because only a small proportion of nucleotide sites are under strong selection (Gossmann et al., 2010; Buffalo, 2021; Dukler et al., 2022; Galtier, 2024). Detecting any relationship between speciation and molecular change might thus require a more targeted approach.

### Rate variation as a key feature in understanding timing of macroevolution

Our analyses confirm that rates of morphological and molecular evolution vary considerably across lineages, even though they lack a strong association with net diversification. Across the 40 data sets analysed here, the degree of morphological rate variation across lineages vastly exceeds that of molecular rates, a finding that is consistent with those of some recent studies (Davies & Savolainen, 2006; Cooper & Purvis, 2009; Janecka et al., 2012; Van Der Wal et al., 2025; Asar et al., in prep). This disparity in the extent of among-lineage rate variation might partly reflect the different approaches to sampling morphological and molecular data (van den Ende et al. 2023), but also suggests contrasting evolutionary modes at the phenotypic and genomic levels. In general, natural selection is likely to be a stronger driver of morphological evolution than molecular evolution, and this factor tends to produce heterogeneous rates of changes across lineages (Gillespie, 1984). In contrast, rates of molecular evolution are potentially driven by factors that act at a whole-genome scale, such as body size, life-history traits, and geographic range size (Bromham, 2009; Snir et al., 2012; Duchêne et al., 2025). The contrasting levels of among-lineage variation between morphological and molecular evolutionary rates can at least partly explain the lack of evidence for a link between the two (Bromham et al., 2002; Asar, et al., 2023, in prep).

We find that levels of rate variation tend to be similar between genomic markers under strong selection and under weak selection, or slightly higher in markers under weak selection. This comparison could be made for 28 out of the 40 data sets analysed in our study. In some cases, among-lineage rate variation is much higher in coding data than in non-coding data, as might be expected if selection tends to produce rate heterogeneity across lineages (Gillespie, 1984; Tong et al. 2017), but there are also instances of the opposite pattern (Fig. 2). This result is somewhat intriguing because we might expect that markers under strong and weak selection are both correlated with life-history traits, leading to a link between levels of among-lineage rate variation in coding and non-coding regions (Nikolaev et al., 2007; Welch et al., 2008; Rubin et al., 2019). Further research is needed to understand the mechanism underlying this pattern and to achieve a detailed characterisation of among-lineage rate variation across genomic regions and for genes in different functional categories.

**Figure 2.**
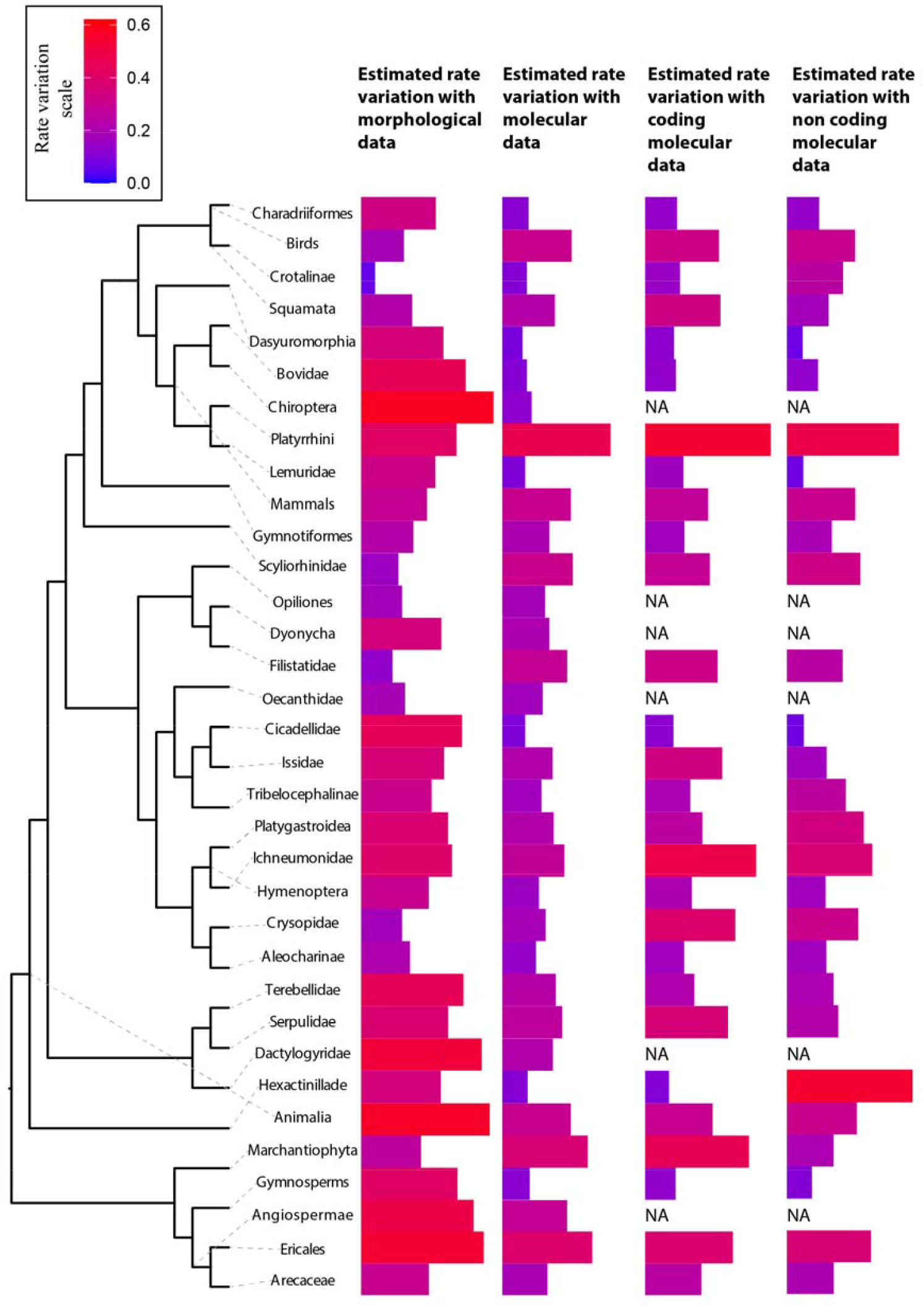
Degree of among-lineage rate variation in genomic regions under strong selection (coding data) and under weak selection (non-coding data), shown for 29 groups of taxa across the eukaryotic Tree of Life. These 29 groups are a sample of the 40 groups analysed in this study.

Our analyses of 40 combined morphological and molecular data sets have revealed marked heterogeneity in evolutionary rates across eukaryotic lineages (Fig. 1), but without compelling evidence of a link between rates of evolutionary change and net diversification. These results are inconsistent with the predictions of the punctuated equilibria model where evolutionary change, either morphological or molecular, is largely tied to speciation. Further analyses will be needed to assess the fit of more complex models of punctuated evolution (i.e., bursts of evolutionary change not necessarily linked to speciation events). Expanding our analyses to a larger data set—incorporating broader taxonomic sampling, additional morphological characters, and more extensive genomic data—will provide deeper insights into the various drivers of phenotypic and genomic change at the macroevolutionary scale. These studies will contribute to a greater understanding of the tempo and mode of evolution across the Tree of Life.

## Methods

### Data

We analysed a collection of 40 data sets that each included a combination of nucleotide sequences and morphological characters. The molecular data sets ranged in size from 2,567 bp to 1,803,947 bp, whereas the morphological data sets ranged from 13 to 3,963 phenotypic characters. Data sets were drawn from various groups of eukaryotes, including Streptophyta, Chordata, Arthropoda, and Annelida. Data sets were modified accordingly for this study. The primary modification involved pruning the phylogenetic tree to include only one representative tip per lineage, in accordance with the taxonomic level under consideration (e.g., one tip per genus). We excluded data sets with taxon sampling at or below the genus level, because these did not permit comparisons of species richness.

For each data set, we identified phylogenetically independent sister lineages using two criteria (Lanfear et al. 2010b). First, the representatives of each lineage needed to form a monophyletic group. Second, each sister-pair had to form a monophyletic group to the exclusion of other taxa in the tree. We excluded any data sets with fewer than four independent sister-pairs.

We partitioned the morphological characters according to their number of possible states. This was done because the inclusion of characters with different numbers of states can lead to underestimation or overestimation of branch lengths if the state space is wrongly assumed to be too large or too small, respectively (Khakurel et al., 2024).

We separated the molecular data into sequences under stronger selection (ribosomal RNA genes, ultraconserved elements, and first and second codon sites of protein-coding genes) and sequences under weaker selection (intergenic regions, introns, and third codon sites of protein-coding genes). This separation was possible for 28 out of the 40 data sets. Following these steps, we produced four separate data matrices for most of the data sets: (i) morphological characters; (ii) nucleotide sequences; (iii) nucleotide sequences under stronger selection; and (iv) nucleotide sequences under weaker selection.

### Branch-length estimation

For each data set, we estimated branch lengths using maximum likelihood in IQ-TREE 2 (Bui et al., 2020). In each analysis, the topology was constrained to match that inferred in the source publication for the data set. The evolution of morphological characters was described using the Mk model, accounting for among-character rate heterogeneity and correcting for ascertainment bias (Mk+G+ASC). For the analyses of molecular data, we aimed to standardize the partitioning scheme across data sets. For alignments composed of partial molecular markers or mitochondrial genomes, we maintained a separate partition for each locus. In the case of genome-scale data, we used the partitioning schemes employed in the source publications. Nucleotide substitution models were selected using ModelFinder (Kalyaanamoorthy et al., 2017).

### Analysis of sister-pairs

For each sister-pair in each data set, we obtained information about the clade size, defined as the number of species described for the order, family, or genus in question. In most cases, this information was extracted from the Catalogue of Life website (Bánki et al., 2024), which consolidates information from various specialized databases and incorporates the contributions of expert taxonomists. Where clade size information was unavailable in the Catalogue of Life, we made use of other databases or specialized literature. We included only sister-pairs that exhibited differences in clade size. In addition, we discarded sister-pairs with short phylogenetic branches (lengths less than the reciprocal of the number of characters). Owing to missing data in some of the morphological and molecular data matrices and to our exclusion criteria, there is some variation in the number of sister-pairs among the various phylogenetic trees estimated for each data set.

### Statistics

To test for a link between the amount of evolutionary change and the diversification process, we used an approach based on phylogenetically independent contrasts or sister-pairs. By definition, the two lineages in each sister-pair originated at the same time, such that they have had equal amounts of time to diversify and accumulate evolutionary change. Therefore, any difference in their morphological or molecular branch lengths reflects a disparity in their respective rates of evolution. Similarly, differences in species richness between the two lineages in each sister-pair reflect differences in net diversification rates.

For each sister-pair, we calculated the differences in: a) morphological branch lengths; b) global molecular branch lengths; c) branch lengths estimated from sites under strong selection; d) branch lengths estimated from sites under weak selection; and e) clade size (species richness). Differences were calculated to ensure that the difference in species richness was consistently positive (Lanfear et al. 2010b). To ensure the phylogenetic independence of the contrasts and to check the assumption of the linear regression that all points have equal variance, we tested whether the absolute mean value of a trait for a sister-pair is related to the size of the difference of that sister-pair (Freckleton 2000). A significant relationship means that the amount of evolutionary change in the trait is a consequence of the shared evolutionary history of the lineages, i.e., the ancestral state of the trait. In addition to the Freckleton test, we followed the recommendation of Garland et al. (1992) to standardize the independent contrasts to ensure homogeneity of variance among the data points. However, unlike Lanfear et al. (2010b), we standardized all branch lengths by the fifth root of the sum of the branch lengths, and clade sizes by the tenth root.

We then used linear regressions through the origin to test for associations between the difference in the amount of evolutionary change and the difference in species richness, following methodology outlined previously (Lanfear et al. 2010a). For each comparison, we considered a correlation to be significant if *p* < α′. The value of α′ was calculated according to the Holm-Bonferroni method for correcting for multiple tests. Since the first comparison of the smallest *p*-value (difference in coding molecular change *versus* species richness in the mammal dataset) did not pass the test (*p-value*= 0.006 > 0.05/136), we stopped the procedure because the testing procedure stops once a failure to reject occurs. Finally, to quantify evolutionary rate heterogeneity among lineages, we estimated the coefficient of variation of root-to-tip distances for morphological trees and for the molecular trees inferred from the three data classes analysed in this study (complete data, regions under strong selection, and regions under weak selection).

